# Worldwide late-Quaternary population declines in extant megafauna are due to *Homo sapiens* rather than climate

**DOI:** 10.1101/2022.08.13.503826

**Authors:** Juraj Bergman, Rasmus Ø. Pedersen, Erick J. Lundgren, Rhys T. Lemoine, Sophie Monsarrat, Mikkel H. Schierup, Jens-Christian Svenning

**Affiliations:** Center for Biodiversity Dynamics in a Changing World (BIOCHANGE) & Section for Ecoinformatics and Biodiversity, Department of Biology, Aarhus University, DK-8000 Aarhus C, Denmark; Bioinformatics Research Centre, Aarhus University, DK-8000 Aarhus C, Denmark

## Abstract

The worldwide loss of large animal species over the past 100,000 years is evident from the fossil record, with climate and human impact as the most likely causes of megafauna extinctions. To help distinguish between these two scenarios, we analysed whole-genome sequence data of 142 species to infer their population size histories during the Quaternary. We modelled differences in population dynamics among species using ecological factors, paleoclimate and human presence as covariates. We report a significant population decline towards the present time in more than 90% of species, with larger megafauna experiencing the strongest decline. We find that population decline became ubiquitous approximately 100,000 years ago, with the majority of species experiencing their lowest population sizes during this period. We assessed the relative impact of climate fluctuations and human presence on megafauna dynamics and found that climate has limited explanatory power for late-Quaternary shifts in megafauna population sizes, which are largely explained by *Homo sapiens* arrival times. As a consequence of megafauna decline, total biomass and metabolic input provided by these species has drastically reduced to less than 25% compared to 100,000 years ago. These observations imply that the worldwide expansion of *H. sapiens* caused a major restructuring of ecosystems at global scale.

## Introduction

The late-Quaternary extinction event^1,2^ is characterised by selective extinctions of large-bodied animals (megafauna) at a global scale. At the present date, a small fraction of the historically speciose megafaunal groups persist in rapidly diminishing communities, many of which face immediate threat of extinction^3,4^. The causes of megafauna decline have been subject to long-standing debate, with paleoclimate fluctuations and human-related activities emerging as the main explanatory factors^5–17^.

Studies of past megafauna dynamics focus on fossil data to infer changes in species distributions and extinction rates. While the fossil record provides valuable insight into species’ histories, the fragmentary nature of this data results in a limited resolution of past population dynamics. Additionally, past population sizes are difficult to determine from such data. However, inference of past population size dynamics using DNA sequence data is an established approach in genomic studies^18–20^. These approaches are generally based on the sequentially Markovian coalescent (SMC) framework^21^, allowing inference of times to the most recent common ancestor at every nucleotide site along the genome. This information can in turn be used to reconstruct the time-resolved trajectory for a species’ population history from up to a million or more years in the past^22,23^. Given the requirement of genome-wide determination of nucleotide diversity, SMC-based methods initially focused on the inference of human population size fluctuations^18^. However, the rapid increase in cost-effectiveness and quality of next generation sequencing (NGS) technologies^24–27^ now allows for these methods to be applied to a wide variety of animal and plant species.

High-quality reference genome assemblies and short read sequencing data have now become available for a large fraction of terrestrial mammal megafauna. Consequently, studies of SMC-derived histories across entire mammalian clades are becoming increasingly common^28–38^. These studies have the potential to provide a complementary view to canonical fossil-based studies of extinctions by ascertaining the driving factors of past population size dynamics in extant megafauna, but a global overview of megafauna SMC histories in the context of past climatic shifts and human impact is lacking.

In this study, we curate a dataset of DNA sequence data for 142 extant terrestrial megafauna mammals and implement a common bioinformatic pipeline for inference of SMC-based population histories. We study population dynamics of megafauna as a function of species’ ecology, geographical distribution, climate and anthropogenic influence. We detect a global, severe decline in megafauna population sizes over the past 100,000 years. These observations are best explained by the influence of human worldwide expansion rather than past climatic conditions.

## Results

### General decline of megafauna populations throughout the Quaternary

We used a common bioinformatic pipeline to infer past population size changes of extant megafauna from curated diploid genome sequences (Table S1), with the time frame of population size estimates covering the Quaternary period (2.58 million years ago until present) for the majority of studied species. The pairwise sequential Markovian coalescent (PSMC)^18^ curves in Figure 1A summarise population dynamics estimated from 142 megafauna genomes, separated by ecological realm (Figure 1B). We estimate that the most severe decline in population size occured in the Nilgiri tahr (*Nilgiritragus hylocrius*) with the 95% highest posterior density interval (HPDI) for the slope of population size change in the range [-0.708, -0.467], while the springbok (*Antidorcas marsupialis*) experienced the strongest, yet non-significant, increasing population trend (95% HPDI: [-0.125, 0.153]). Generally, megafauna population sizes decrease towards present time, with the 95% HPDI for the slope of population size change significantly below zero for 91% (129/142) of the species and a negative mean slope for 99% (141/142) of the species.

**Figure 1.**
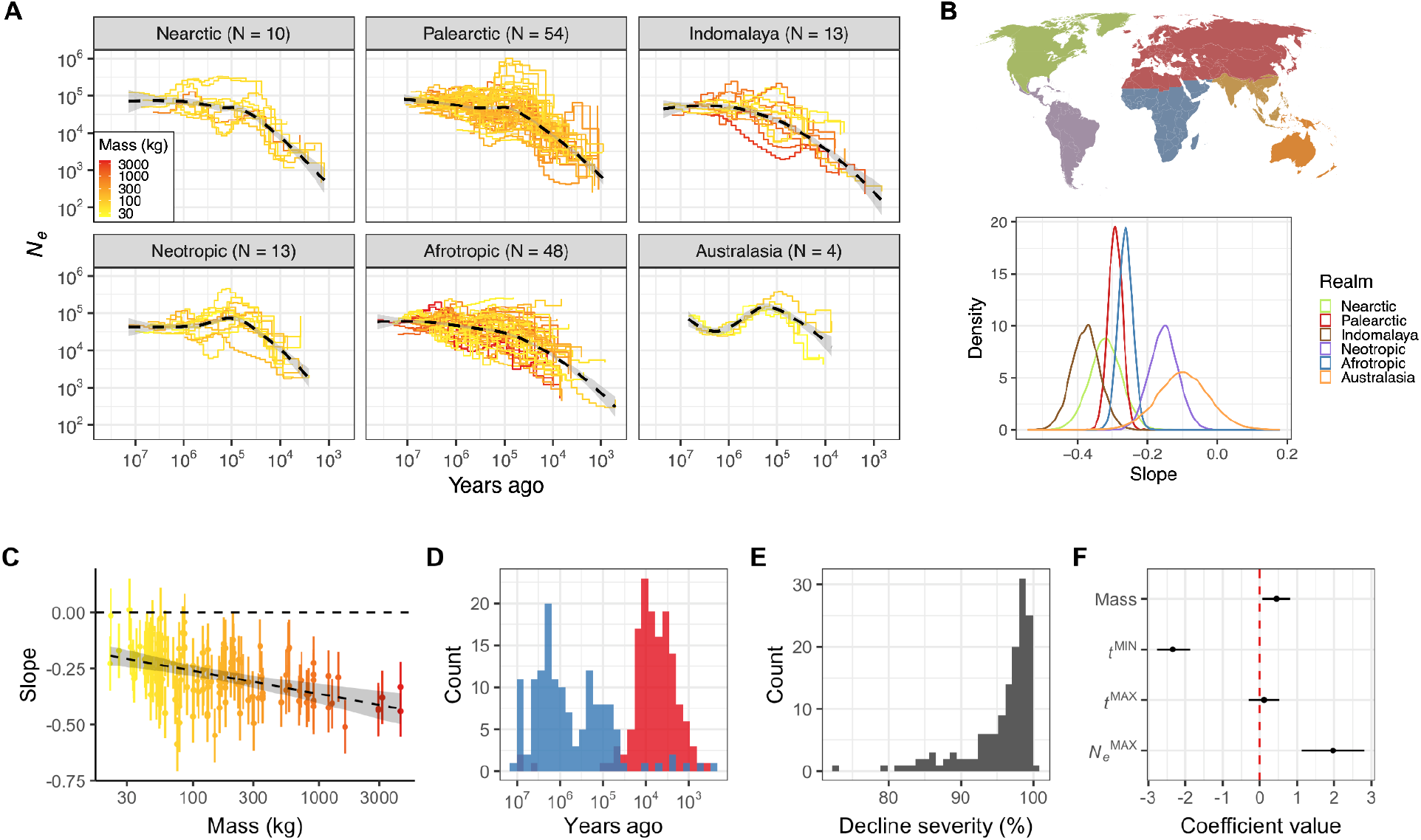
Population size (*N*_*e*_) dynamics of 142 extant megafauna species.**A**. Each step line represents changes in *N*_*e*_ with respect to time (in years) for a single megafauna species, colored by a gradient based on average adult mass. Panels separate species by ecological realms. The number of species within each realm is provided in parentheses. The dashed lines are average loess regression lines for the relationship between *N*_*e*_ and time within a realm. Both axes are log_10_-transformed. **B**. Slope of population size change given division of species with respect to their biogeographic realm. **C**. Relationship between species’ adult mass and the slope of population size change. The x-axis is log_10_-transformed. **D**. Distribution of times since species experienced their highest population size (blue) and lowest population size (red). **E**. Distribution of species’ decline severity. **F**. Coefficient values of explanatory variables (species’ adult mass; *t*^MIN^: time since a species achieved the lowest population size, *t*^MAX^: time since a species achieved the highest population size, *N*_*e*_ ^MAX^: highest population size achieved during the whole time span) for a regression model with species’ decline severity as the response variable. The distribution for each coefficient is the 95% HPDI, with the point representing the median. The red dashed line represents no effect.

Population declines varied across ecological realms (Figure 1A, B), with Australasia and the Neotropics experiencing the least severe declines over the Quaternary period (95% HPDI: [-0.244, 0.044] and [-0.228, -0.070], respectively), compared to Indomalaya and Nearctic (95% HPDI: [-0.458, -0.299] and [-0.410, -0.227], respectively). Separation of species according to the biome they occupy (Figure S1A) resulted in the largest discrepancy of population size decline between polar (95% HPDI: [-0.317, 0.037]) and temperate-adapted species (95% HPDI: [-0.460, -0.296]), while insectivores (95% HPDI: [-0.228, 0.201]) experienced a small and non-significant decrease compared to hypercarnivores (95% HPDI: [-0.394, -0.223]; Figure S1B).

Lastly, species with ranges overlapping regions where *Homo sapiens* was the first and only hominin present, tend to have the lowest decline (95% HPDI: [-0.269, -0.155]), compared to species in regions where archaic *Homo* species arrived early (95% HPDI: [-0.380, -0.290]; Figure S1C). Generally, non-African temperate regions with a relatively long history of hominin activity experienced the largest decrease in megafauna population sizes. In contrast, and with the exception of polar species, warmer biomes with only *H. sapiens* activity seem to have declined the least. However, this observation is most likely driven by an increase of megafauna population sizes in Neotropics and Australasia between 1.25 million and 100,000 years ago, prior to human arrival (Figure 1A). Notably, population decline starting at approximately 100,000 years ago, and continuing towards the present, is ubiquitous across realms.

To test if the size selection bias observed for recently extinct species^39^ is reflected in extant megafauna population dynamics, we considered the relationship between the slope of population size change and species’ adult mass. We observed a significantly negative relationship between mass and slope (95% HPDI: [-0.152, -0.059]), indicating that larger species experienced stronger declines during the Quaternary (Figure 1C). The majority of species experienced the lowest population sizes closer to present time compared to their highest past sizes (*t =* -7.777, *p* < 0.001; Figure 1D). Furthermore, the severity of decline, defined as the percentage of population size decrease with respect to the highest past population size, was exceptionally high, with 95% of species experiencing a population decline between 84.3% and 99.9% (Figure 1E).

We then modelled decline severity as a function of species’ mass, time since species experienced the highest and lowest population sizes, and the maximum population size achieved in a history of a species (Figure 1F). We observed a significantly positive relationship between decline severity and species’ mass, in line with the observed size selection bias (Figure 1C). We also found a positive relationship for maximum population size, suggesting that the population decline in species with larger past population sizes has been more severe during the Quaternary, or that species with lower population sizes experienced a milder population decline, indicative of an increased potential for decline in species with large population sizes. Lastly, we observed a strong negative relationship between decline severity and time since a species experienced a lowest past population size. This observation demonstrates that although the majority of lowest megafauna population sizes are observed close to present time (Figure 1D), variation within these times is still informative for severity of decline. Moreover, this negative relationship shows that population declines have become increasingly more extreme towards the present time.

### Climate-based models are unable to predict population decline during the last 100,000 years

To better understand the recent population decline of megafauna, we focused our analysis on population trends during the last 742,419 years (Figure 2A) for which we have high-quality estimates of global temperature change^40^. Specifically, we were interested in whether past climatic conditions predict the recent severe declines in megafauna population sizes. We therefore divided the inferred population sizes across species into two time-dependent categories, before and after 100,000 years ago. This time-point was chosen as it includes the time range during which population decline intensified - between 92,044 and 115,878 years ago, as estimated by breakpoint analysis - and it facilitates subsequent division of the last 100,000-year period into discrete time windows. We used all estimates of population sizes between 100,000 and 742,419 years ago to fit a model with climatic predictors, while population size estimates younger than 100,000 years were predicted using the best-fitting model and compared to the observed values. As predictors, we used the average temperature of the focal time window for which we have an estimate of a species’ population size, as well as average temperature of the preceding time window (*i*.*e*., temperature lag). Model fitting and prediction were conducted separately for each species.

**Figure 2.**
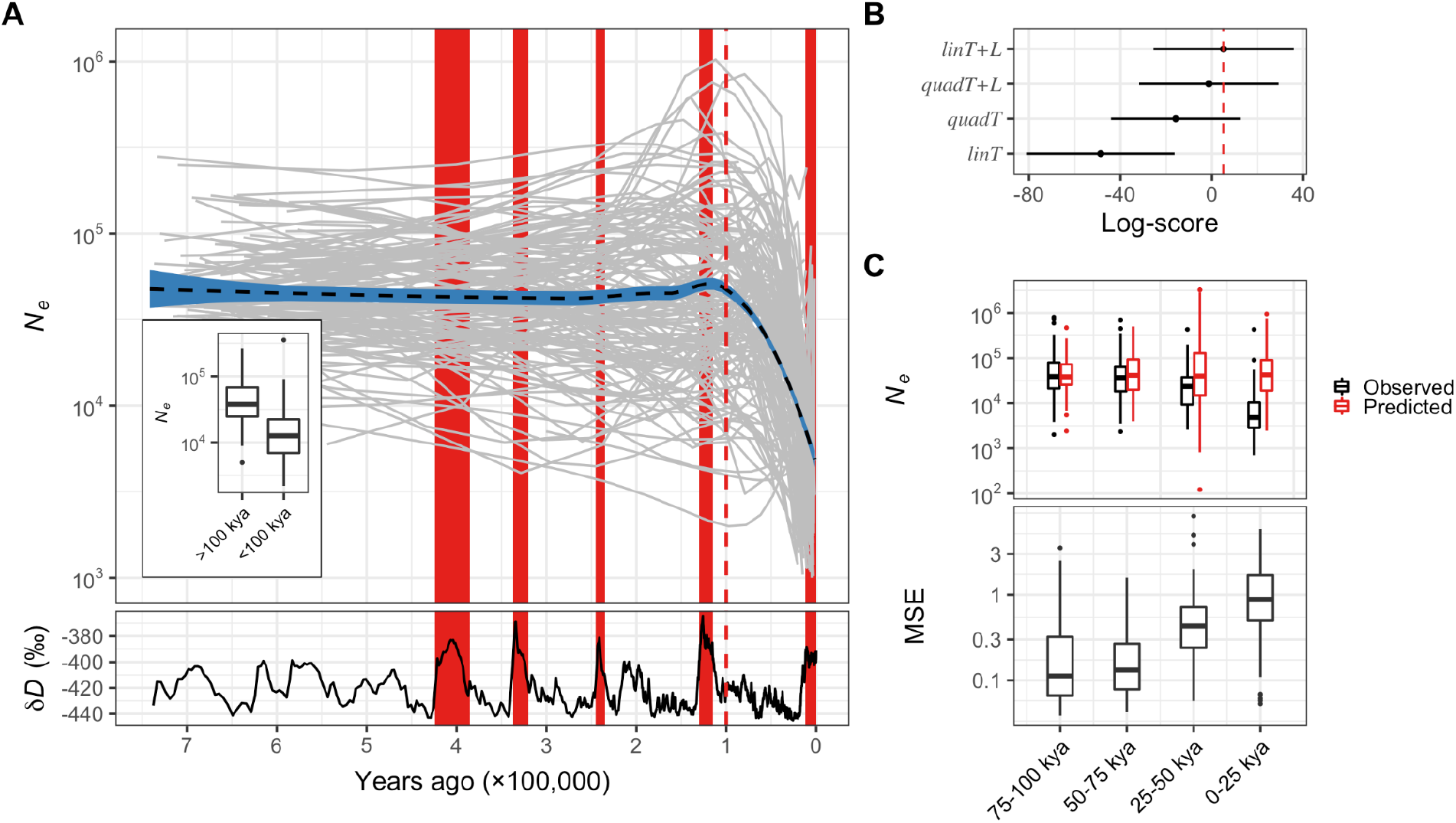
Climate-based models of population size (*N*_*e*_) change over the last 742,419 years.**A**. Each grey line in the top panel represents an *N*_*e*_ trajectory (log_10_-transformed) with respect to time (in years) for a single megafauna species. The inset shows the distribution of average population sizes for each species in the time periods prior and after 100,000 years ago (vertical red dashed line). The black dashed line with the blue ranges is the average population size trend across all species. The bottom panel is past temperature change with warming periods highlighted in red. **B**. Log-score of leave-one-out cross-validation for climate-based models, with the red dashed line indicating the best-fitting model. **C**. Distributions of population size (top panel) and mean squared errors (MSE; bottom panel) of the best-fitting model, for four time intervals during the last 100,000 years. Both y-axes are log_10_-transformed.

We fitted two models of the dependence between population size and climate. First, we assumed a linear relationship between the population size response and the climate explanatory variables. Secondly, we modelled a quadratic relationship between the variables, thus assuming that a species may experience the highest population size at some optimal temperature value, with a decline in size above or below the temperature optimum. We fit both models with and without the inclusion of the temperature lag predictor and use leave-one-out cross-validation to evaluate the predictive accuracy of the models (Figure 2B). We find that the model with only a linear effect of average temperature has the lowest log-score from the cross-validation scheme, and thus lowest predictive accuracy. On the other hand, the quadratic models and the linear model with both the temperature and lag predictor have higher log-scores and similar predictive accuracy.

Finally, we use the best-fitting model (with linear temperature and lag predictors) to predict the population sizes of megafauna during the last 100,000 years. Figure 2C shows the relationship between the observed and predicted population sizes for four consecutive 25,000-year time windows over the last 100,000 years. Notably, the difference between the observed and predicted values is larger for time windows that are closer to the present. The difference is non-significant for the oldest time window between 75,000 and 100,000 years ago (*t =* -0.028, *p* = 0.977), but gets progressively more significant closer to the present (50,000-75,000 years ago: *t =* -1.716, *p* = 0.087; 25,000-50,000 years ago: *t =* -4.436, *p* < 0.001; 0-25,000 years ago: *t =* -15.726, *p* < 0.001). This trend is also reflected in the increasing mean squared error (MSE) of the model fit. In conclusion, we detect a time-dependency of model performance indicating the inability of climate changes to predict population size shifts over the past 75,000 years. Therefore, additional factors must be taken into consideration when modelling megafauna dynamics. In the next section, we focus on anthropogenic predictors.

### Models with human impact accurately capture recent population size dynamics

Here, we were interested in assessing the relative impact of climate and anthropogenic predictors on past megafauna dynamics. We use the full dataset of population size estimates to fit the models and assess explanatory power for every predictor combination. The basic model type only includes climate predictors from the previous section. For the second model type, we introduce human impact predictors as a function of *Homo sapiens* arrival times to each ecological realm^16^. We consider two arrival-informed models. Firstly, we consider the overlap between the human arrival range and a focal time window for which we have an estimate of a species’ population size. The extent of the overlap determines the probability of human presence within the geographic range of the species, *i*.*e*., the likelihood of human-megafauna interaction during a specific time period. We use this probability as a predictor in a linear model with population size as the response variable. However, probability of human presence is likely a conservative proxy for human impact, as the influence of *H. sapiens* likely continued to increase post-arrival. We therefore introduce a second arrival-informed model where population size is assumed to be constant prior to human arrival, after which population size follows either a linear or a non-linear (logistic or exponential) trend. Both arrival-informed models are combined with climate-informed predictors to construct the third model type which incorporates the joint effect of humans and climate on megafauna dynamics. In total, we consider 24 models (Table S6) with climate only (4 models), human only (4 models) or combined predictors (16 models), and use leave-one-out cross-validation to compare model performance (Figure S2).

Notably, models that include non-linear population size change after human arrival have the highest predictive accuracy (Figure 3A). The model with the overall highest predictive accuracy includes only human arrival time as a predictor and assumes a logistic trend of megafauna population change post-arrival. The 95% HPDI for the rate of change of the logistic trend is below zero for 68% (97/142) of the species, with a median negative rate for all species, indicating that the majority of megafauna species experienced a gradually accelerating decline in population size after human arrival. This is consistent with cumulative human impacts on megafauna populations post-arrival, as a consequence of gradual establishment of human populations in a region. Correspondingly, the best-fitting models with combined predictors again include the logistic trend of population change post-arrival. The human only model with an exponential population size change is nested within models with combined predictors, likely indicating that the exponential phase of the logistic trend dominates megafauna dynamics post human arrival. The second best-fitting models are human only models that contain a linear relationship between population size and human predictors. Lastly, the four climate only models have lowest log-scores and therefore poorest predictive accuracy.

**Figure 3.**
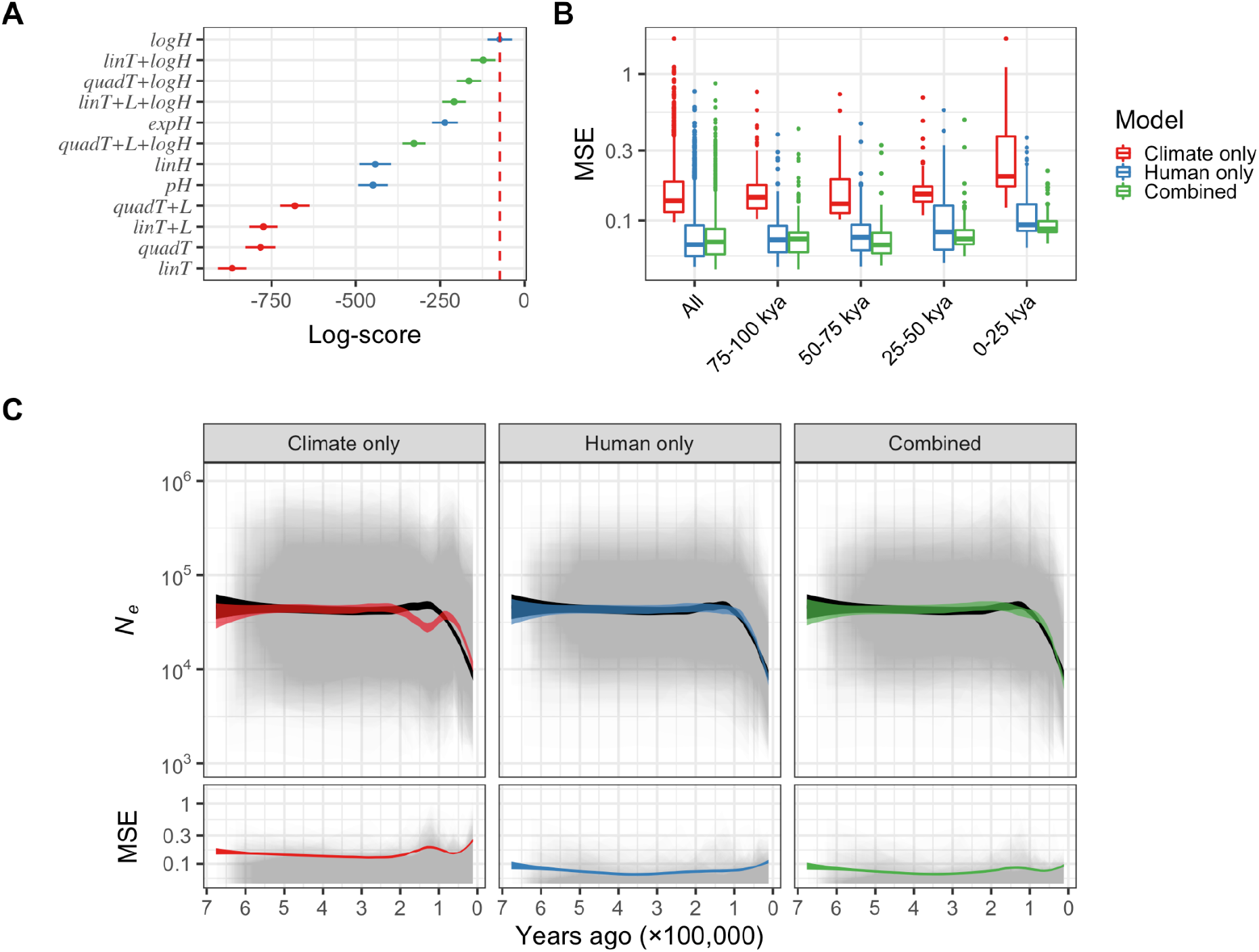
Climate and human arrival-informed models of population size (*N*_*e*_) change over the last 742,419 years.**A**. Log-scores of leave-one-out cross-validation for all climate and human only models, and four best-fitting models with combined predictors. The red dashed line indicates the best-fitting model. **B**. Distributions of mean squared errors (MSE) for the best-fitting model in each model class, for the whole time span (All: 0-742,419 years ago) or four time intervals during the last 100,000 years. A single MSE value corresponds to one specific model and species. The x-axis is log_10_-transformed. **C**. The top panels show the mean observed population size trend of megafauna and per-species posterior predictive distributions of the best-fitting model in each model class. The solid black area is the mean observed population size trend, while the overlapping gray areas represent the 95% highest posterior density interval (HPDI) ranges for all species and the colored areas are mean fitted population size trends based on mean posterior values. The bottom panels show the corresponding MSE values. Both y-axes are log_10_-transformed.

In Figure 3B we show the distribution of per-species mean MSE values for the best-fitting model in each class. Across the whole time span, the climate only model has a significantly higher MSE compared to the human only (*t =* 24.225, *p* < 0.001) and combined models (*t =* 25.438, *p* < 0.001), while the difference between MSE distributions of the human only and combined models is non-significant (*t =* 1.517, *p* = 0.129). A similar trend is observed for each of the four discrete time windows during the last 100,000 years. Furthermore, an increase in MSE closer to present time is observed for all model classes. However, the difference between MSE distributions of the two most recent time intervals is significant for the climate only (*t =* -6.953, *p* < 0.001) and combined models (*t =* -2.305, *p* = 0.022), and non-significant for the human only model (*t =* -0.849, *p* = 0.397), reflecting the time-dependency of model performance for models with climatic predictors. Notably, the posterior predictive distributions across the past 742,419 years show that the largest discrepancies between the observed and predicted population sizes are present for time windows around the Last Interglacial (Eemian) period (115,000-130,000 years ago), especially for the climate only model (Figure 3C). This is likely caused by many species experiencing large population sizes during the Eemian interglacial (Figure 2A, Table S1), which had similar climatic conditions as the current warming period (Holocene; < 11,700 years ago), during which populations were generally strongly reduced (Table S1). The climate only model therefore compensates between highest and lowest past population sizes during the last two warming periods by underestimating population sizes for the Eemian period, while overestimating them for the Holocene period. Additionally, the inconsistency in the ranking of best-fitting climate-based models when different time periods are considered (Figure 2B, 3A) again points to the inadequacy of these models in explaining megafauna dynamics. In contrast, models that include human arrival-informed predictors show a greater correspondence between mean observed and predicted population trends, lower variance of posterior predictive distributions and lower MSE across the whole time span (Figure 3C).

### Consequences of megafauna decline

The stark decline of megafauna populations during the last 100,000 years is expected to drastically change ecosystem composition and functioning^41^. To estimate the magnitude of these effects, we calculated an average baseline for total effective population size, biomass, and metabolic rate contributed by megafauna during the period prior to 100,000 years ago, and compared it to time periods that are closer to the present (Figure 4A). Generally, total megafauna population size and metabolic rate were highest between 75,000-100,000 years ago, and even exceeded the baseline average size until 50,000 years ago. Similarly, total megafauna biomass was higher than the corresponding baseline values until 75,000 years ago. These observations can be explained by an expansion of megafauna during, and immediately following, the Last Interglacial period (Figure 2A, Table S1). However, all three parameters show a continuous decline towards the present, finally reducing to less than 25% of their average baseline values in the youngest timeframe (0-25,000 years ago). Notably, total biomass had a larger decline compared to total population size, again illustrating the size selection bias in decline dynamics (Figure 1C, F).

**Figure 4.**
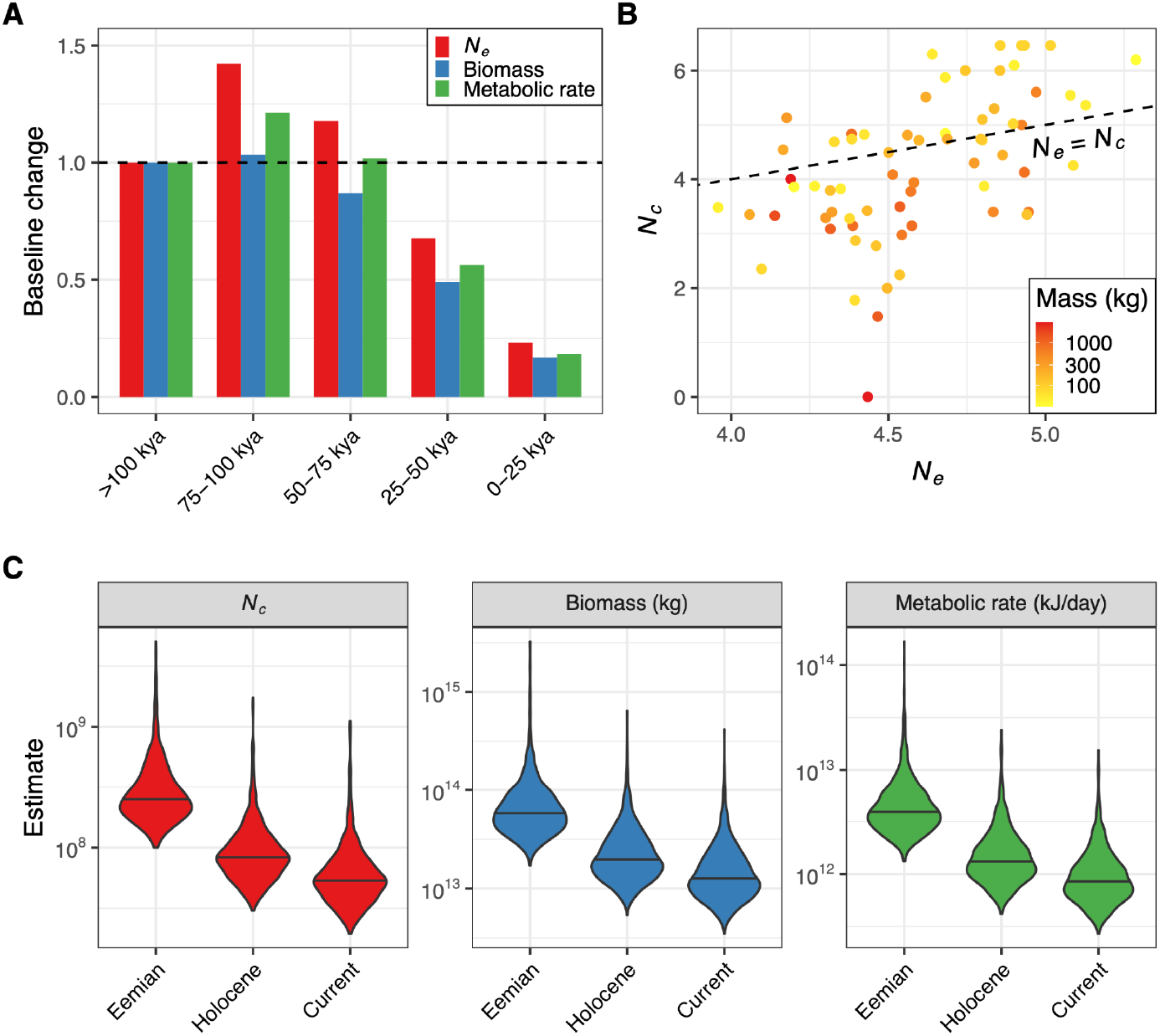
Consequences of recent megafauna decline.**A**. Change in total effective population size, biomass and metabolic rate relative to the average baseline values calculated for the period between 100,000 to 742,719 years ago (dashed line). **B**. Relationship between average effective size (*N*_*e*_) calculated for the period between 100,000 to 742,719 years ago and present-day census size (*N*_*c*_) for 66 megafauna species. Both axes are log_10_-transformed. The dashed line is the 1:1 line. **C**. Posterior sample distributions of the sum across all species for total megafauna census size, biomass and metabolic rate for the Eemian (115,000-130,000 years ago), Holocene (< 11,700 years ago) and current period. Each distribution consists of 1,000 posterior samples.

We next explore the relationship between the average effective population sizes prior to 100,000 years ago, and current census sizes (*N*_*c*_) of megafauna estimated by IUCN. Strikingly, out of 67 species from our dataset for which census size estimates are available, 58% (39/67) have a lower census size compared to the past effective size (Figure 4B), indicative of recent and strong population bottlenecks in these species^42,43^. Furthermore, these species tend to have higher adult mass (*t =* -3.003, *p* = 0.004), again signifying stronger population declines experienced by larger megafauna.

A decrease in census population size of a species is expected to ultimately result in a decrease in effective population size. Therefore, the two population size measures are expected to track each other, as demonstrated by their positive correlation (Figure 4B; Spearman’s ϱ = 0.558, *p* < 0.001). This relationship can be utilised to predict megafauna abundance during different periods of time. Specifically, we are interested in comparing megafauna census sizes between the Eemian and Holocene periods, given their similarity in climatic conditions. To achieve this, we fit a linear model for the relationship between *N*_*e*_ values estimated for the Holocene period and IUCN *N*_*c*_ estimates, while only considering species for which the Holocene effective size does not exceed current census size (*i*.*e*., *N*_*e*_/*N*_*c*_ < 1; Figure S4). In effect, this model predicts megafauna census sizes that might be expected in the absence of severe bottlenecks. We then generate posterior sample distributions for the total sum of megafauna census size, biomass and metabolic rate across all species for both the Eemian and Holocene period (Figure 4C). Additionally, we estimate posterior sample distributions for current time, by scaling the Holocene estimates with a factor that takes into account the average decline severity experienced by megafauna. On average, this factor scales down current census sizes to ∼64% of the estimated Holocene census sizes.

We estimate that the mean census size summed across the 142 species studied was 345 (median of 248) million individuals during the Eemian period, with Holocene and current estimates of 108 (median of 84) and 70 (median of 54) million individuals, respectively. The estimates have large variances with 95% HPDIs of [100, 761], [30, 235] and [20, 152] million individuals for the Eemian, Holocene and current periods, respectively. On average, the Eemian period was host to ∼3-5× more megafauna individuals compared to the Holocene, with a similar increase in total biomass and metabolic rate output. These results indicate that the climatic conditions of the Holocene are likely suitable for accommodating a substantially greater number of large animals than are present in existent ecosystems.

## Discussion

Our results show that over the last 100,000 years, megafauna communities have been severely decimated not just in species numbers through extinctions worldwide, but also through severe reductions in population sizes of the surviving species. Further, analogous to the strong size-selectivity of the extinctions, the population declines were most severe for the bigger species. Our results hereby show that terrestrial mammal faunas worldwide have been even more severely downsized across the late-Quaternary than indicated by the extinctions, representing a major restructuring of ecosystems at a global scale. We also show that this downsizing was unique relative to earlier in the Quaternary and that human presence was the main driving factor, as opposed to fluctuating climatic conditions. The inability of climate to predict the observed population decline of megafauna, especially during the past 75,000 years (Figure 2C), implies that human impact became the main driver of megafauna dynamics around this date. Importantly, given that we use human impact to predict trajectories of effective, rather than census population sizes, we hypothesise that anthropogenic influence on megafauna dynamics is likely underestimated in our models. In line with this proposition, we observe that many megafauna species have higher effective population sizes compared to current census sizes (Figure 4B).

The recent extinctions of a large number of megafauna species^1,2,13^ resulted in multiple co-extinctions and reduction of diversity due to the loss of important ecological roles these species performed across various ecosystems^41^. Although such events might have provided opportunities for population expansion in surviving species through competitive release, the observed decline of extant megafauna during this time indicates that such a scenario was never realised. Moreover, the majority of extant megafauna are currently met with an unprecedented severity of extinction risk, casting further uncertainty on the survival of existing ecosystems. The potential of the current epoch for species restoration can be glimpsed from the patterns of megafauna abundance predicted for the Eemian interglacial (Figure 4C), given its climatological similarity to the Holocene. Importantly, the fulfilment of this potential would require urgent planning at a global scale and reinforcement of current conservation and restoration efforts^3,4^.

## Methods

### Data curation

The latest available reference genome assemblies for each species were downloaded from https://www.ncbi.nlm.nih.gov/. For short read mapping, we chose data from one representative biosample per species, corresponding to the individual used for reference assembly or an individual with the highest amount of short read data available. Temperature data were taken from Augustin *et al*. (2004)^40^.

### Mapping of short read data

The fastq files containing short read data were downloaded from https://www.ncbi.nlm.nih.gov/ and processed by picard tools (https://broadinstitute.github.io/picard/) to generate an unmapped bam file (FastqToSam module) with marked adapter sequences (MarkIlluminaAdapters module). The program bwa mem v0.7.17^44^ was used to map the reads to reference sequences. Only reference contigs that were more than 1,000 base pairs in length were used for mapping of reads. Secondary alignments and duplicates were removed using picard tools MergeBamAlignment and MarkDuplicates modules. When short read data were spread across multiple files, we merged the resulting files into the final bam file using the picard tools MergeSamFiles module. The average coverage of genomic positions was calculated using the samtools depth program^45^.

### Demography inference

For demography inference, we used the pairwise sequentially Markovian coalescent (PSMC) implementation (https://github.com/lh3/psmc)^44^. To account for potential inference biases introduced by genomic regions with low mapping probability, we created a mappability filter for each reference genome using the snpable program (http://lh3lh3.users.sourceforge.net/snpable.shtml), with the 90% stringency criterion as in Palkopoulou *et al*. (2018)^30^. We used the bcftools mpileup and call modules to call genomic variants from the produced bam files, only for sites within the mappable fraction of the reference genome and on contigs at least 100 kb in length. We extracted the consensus fasta sequence from the resulting vcf file using the vcf2fq module of samtools, using only sites with read coverage of at least ⅓ of the average genomic coverage and no more than twice the average coverage for a particular species. The PSMC input was created using the fq2psmcfa tool (-q20), followed by demography inference with three different settings for the -p parameter (“4+25*2+4+6”, “6*1+24*2+4+6” and “10*1+15*2”). We selected a single PSMC output per species that maximises the number of recombination events used to estimate effective population sizes (*N*_*e*_) in each time interval^18^.

Conversion of the PSMC output into effective population sizes and time (measured in years) was done following https://github.com/lh3/psmc^44^. The per generation mutation rate for each species was obtained from literature or predicted using a regression model based on known mutation rates and generation times of extant mammals (Bergeron et al., 2022 unpublished), as described in Supplementary text 1.

### Inference of ecological parameters

Each species was assigned to one ecological realm and biome. To do this, we considered the overlap of the species’ geographic range, estimated using the PHYLACINE database^46^, with each of these geographic classifications. If a species’ range overlapped multiple realms (or biomes), the assignment was conducted by choosing the realm (or biome) with the largest overlap. An analogous procedure was implemented when assigning species to human biogeography regions, which were taken from Sandom *et al*. (2014)^13^. Assignment of species to trophic guilds is based on the corresponding classification from PHYLACINE^46^. Species’ adult mass and metabolic rate were also taken from PHYLACINE. The total biomass and metabolic rates were calculated by multiplying mass and metabolic rate values with population sizes of the corresponding species and then summing the resulting values across all species.

### Statistical modelling

Statistical models used in this study are described in detail in Supplementary text 1. A list of response and explanatory variables used in models, along with their description, is presented in Table S2. All models were fitted using a Bayesian framework implemented in the probabilistic programming package pyMC3 of the Python programming language^47^. All models were run using four Markov chains, each with 2,000 tuning iterations followed by the same number of sampling iterations. Leave-one-out cross-validation of the fitted models was conducted using the Python-implemented ArviZ package^48^.

Breakpoint analysis used to determine the time range during the last 742,419 years for which population size change became more severe was conducted using the “segmented” library^49^ implemented for the R programming language. Both time and population size estimates were log_10_-transformed prior to breakpoint analysis.

## Supplement

### Supplementary Figures

**Figure S1.**
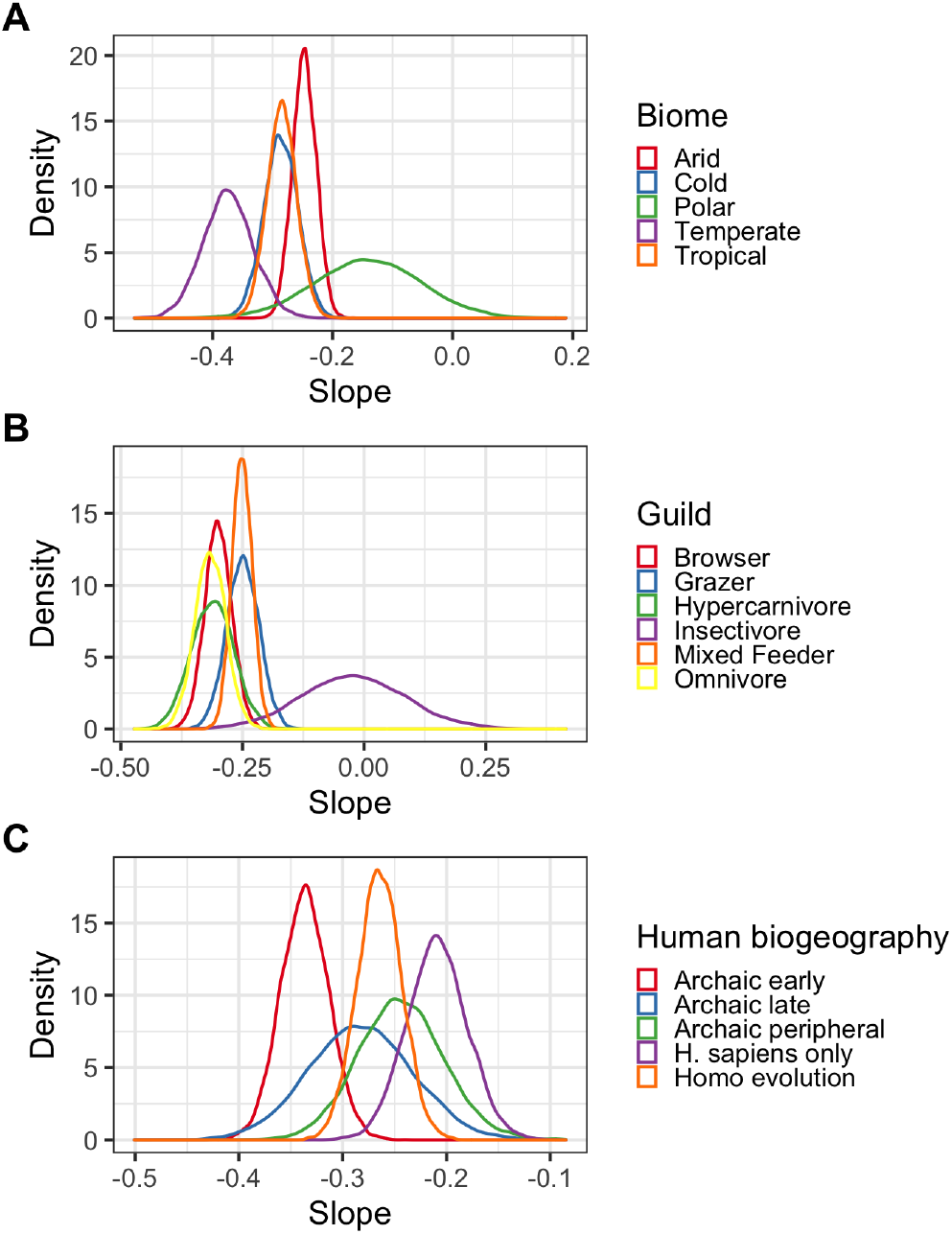
Slope of population size change given division of species with respect to their **A**. biome, **B**. trophic guild and **C**. human biogeography.

**Figure S2.**
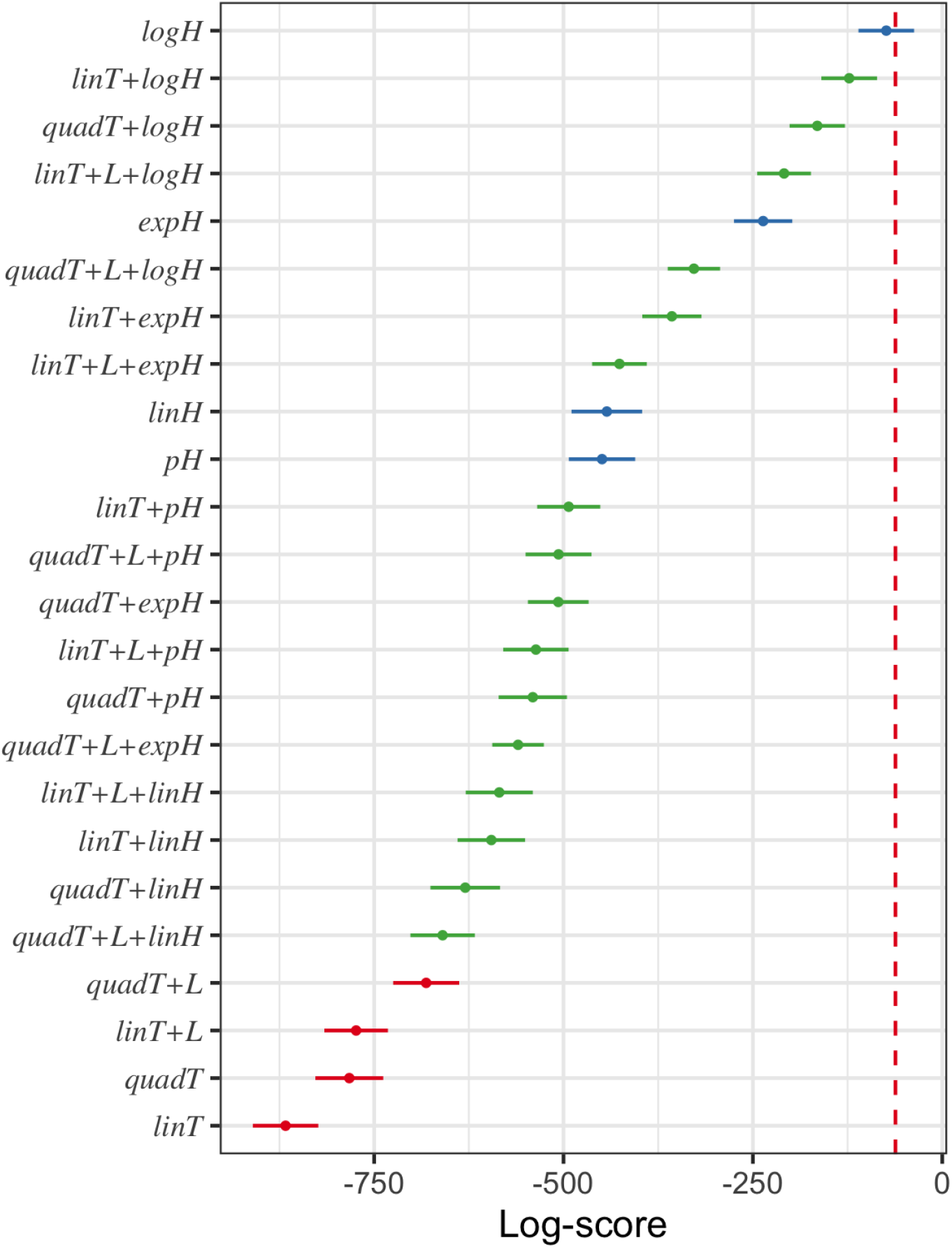
Log-scores of leave-one-out cross validation for all 24 fitted models (Table S6), with the red dashed line indicating the best-fitting model. The colours red, blue and green signify climate only, human only and combined models, respectively.

**Figure S3.**
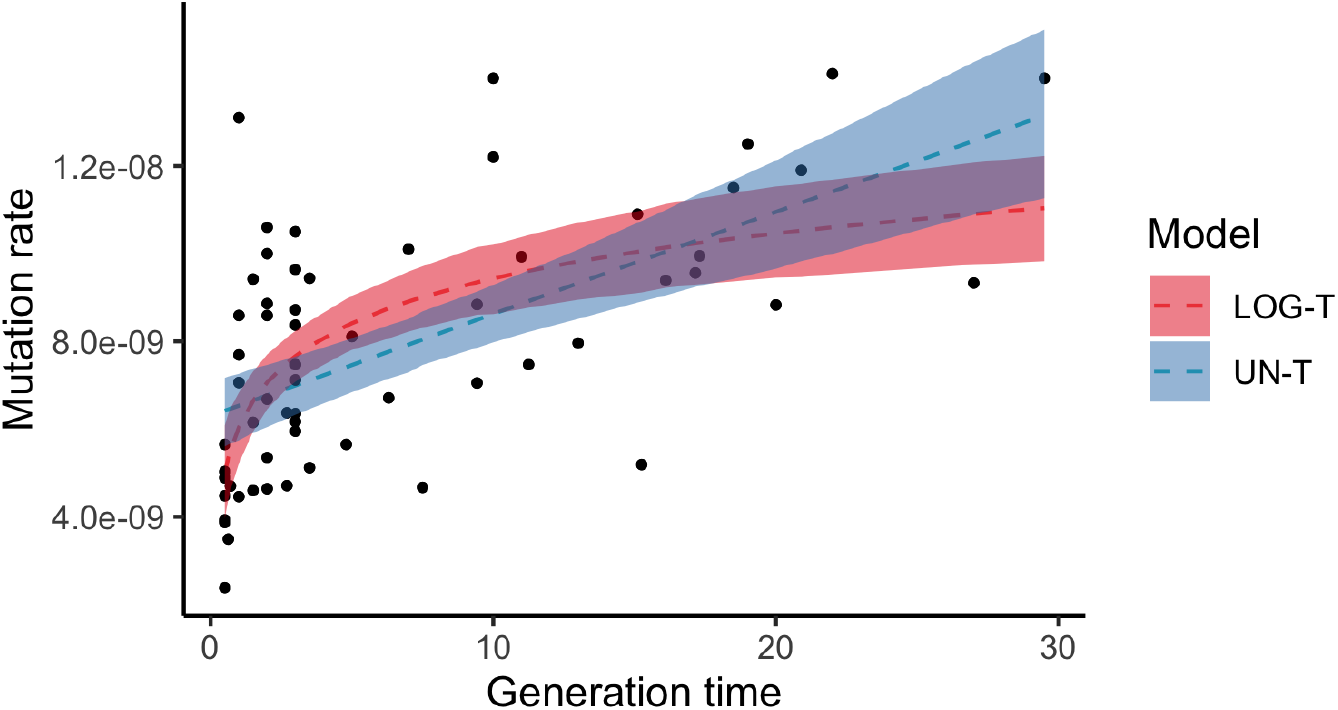
Linear regression model using mammalian trio data from Bergeron et al. (2021, unpublished) with per generation mutation rate as the response variable. The points are the observed data and two different fits depict linear models where the predictor (generation time) was untransformed (UN-T) or log-transformed (LOG-T), respectively.

**Figure S4.**
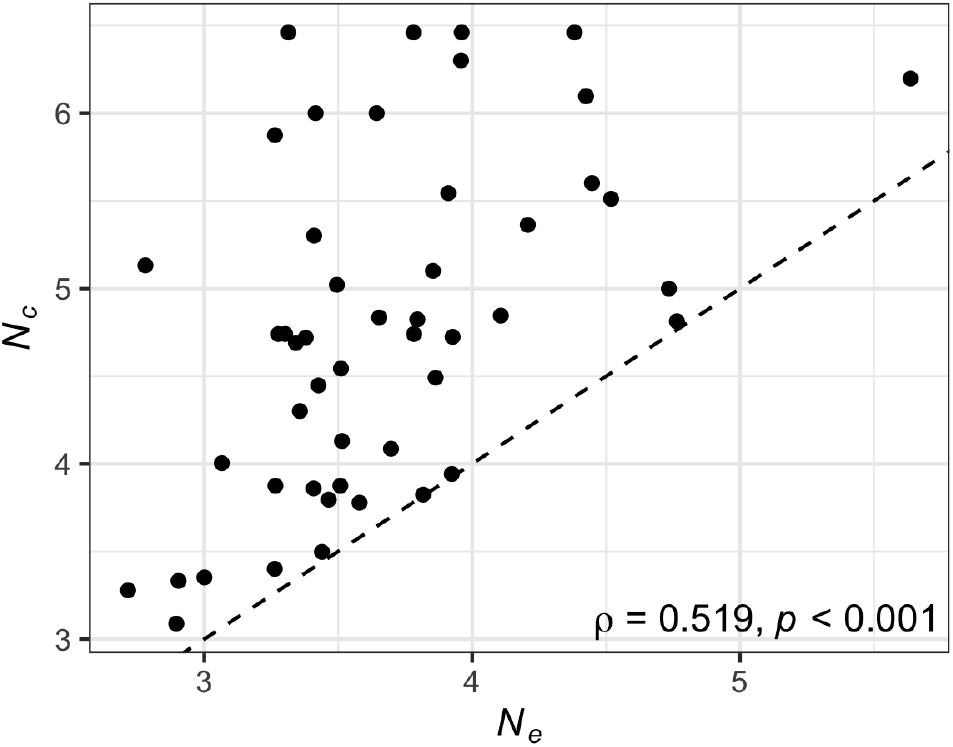
Relationship between Holocene effective size (*N*_*e*_) calculated for the period between 0 to 11,700 years ago and present-day census size (*N*_*c*_) for 49 megafauna species with available IUCN *N*_*c*_ estimates and *N*_*e*_/*N*_*c*_ < 1. Both axes are log_10_-transformed. The dashed line is the 1:1 line with and the Spearman’s ρ correlation value presented in the bottom right corner.

### Supplementary Tables

**Table S1.**
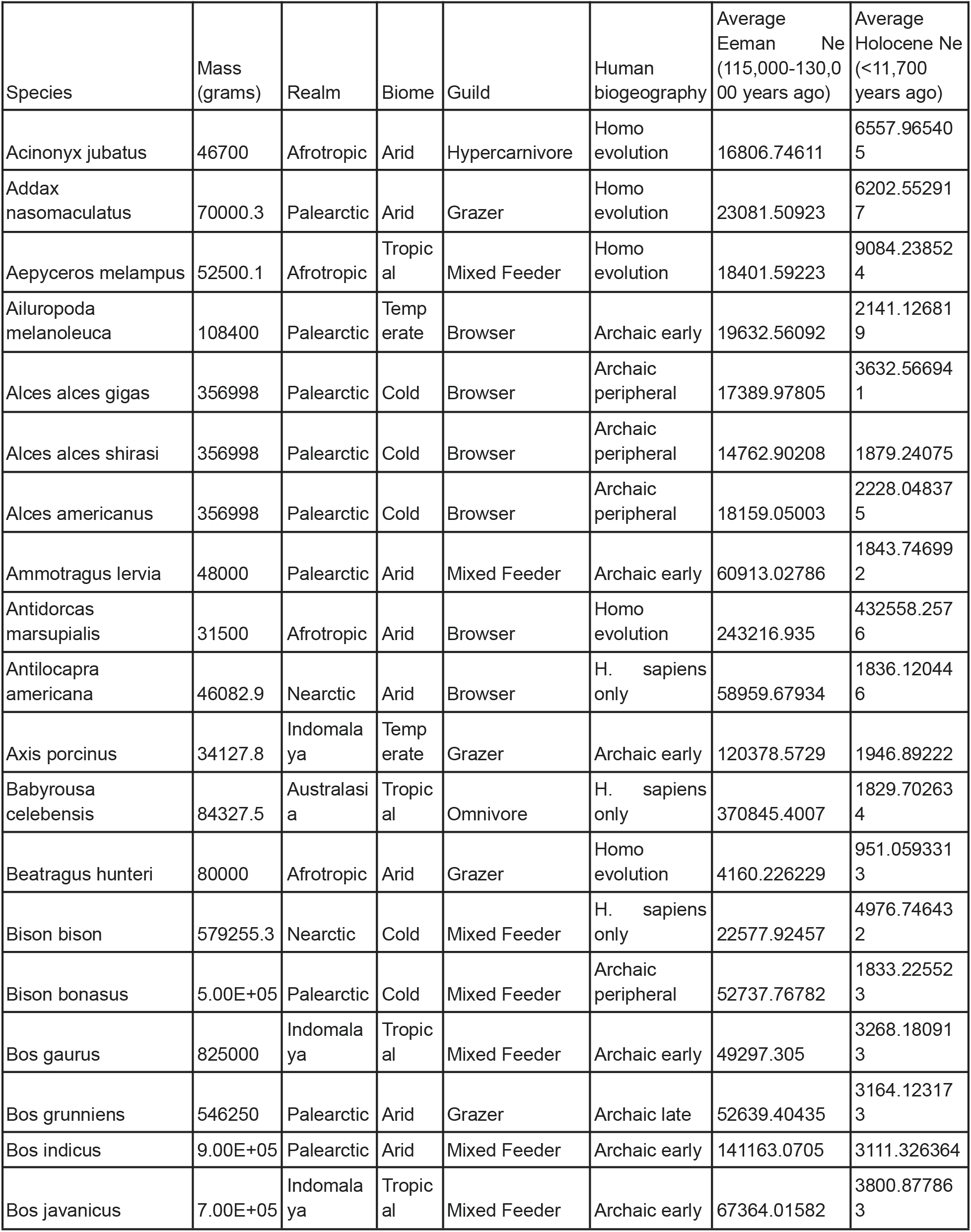
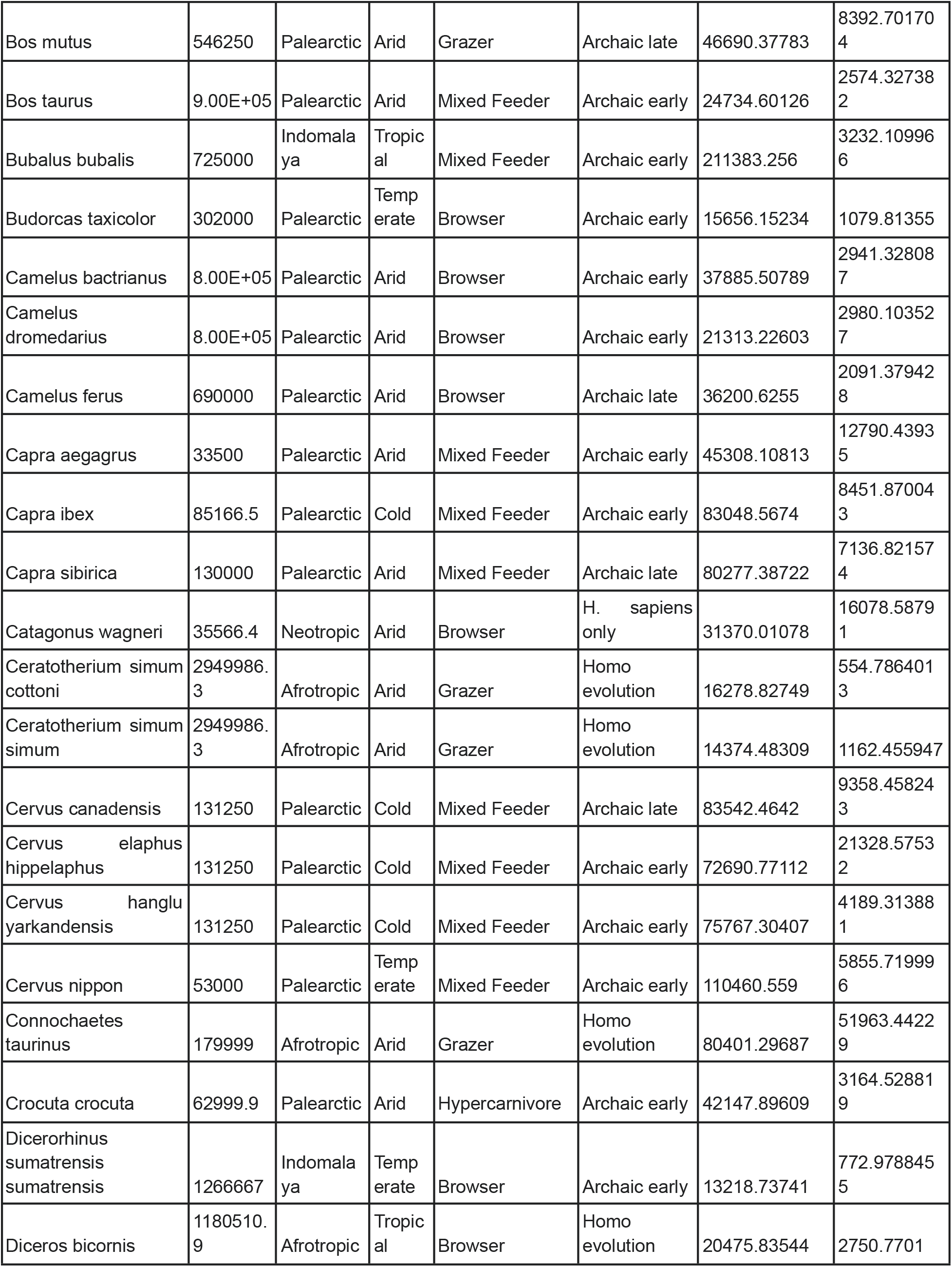
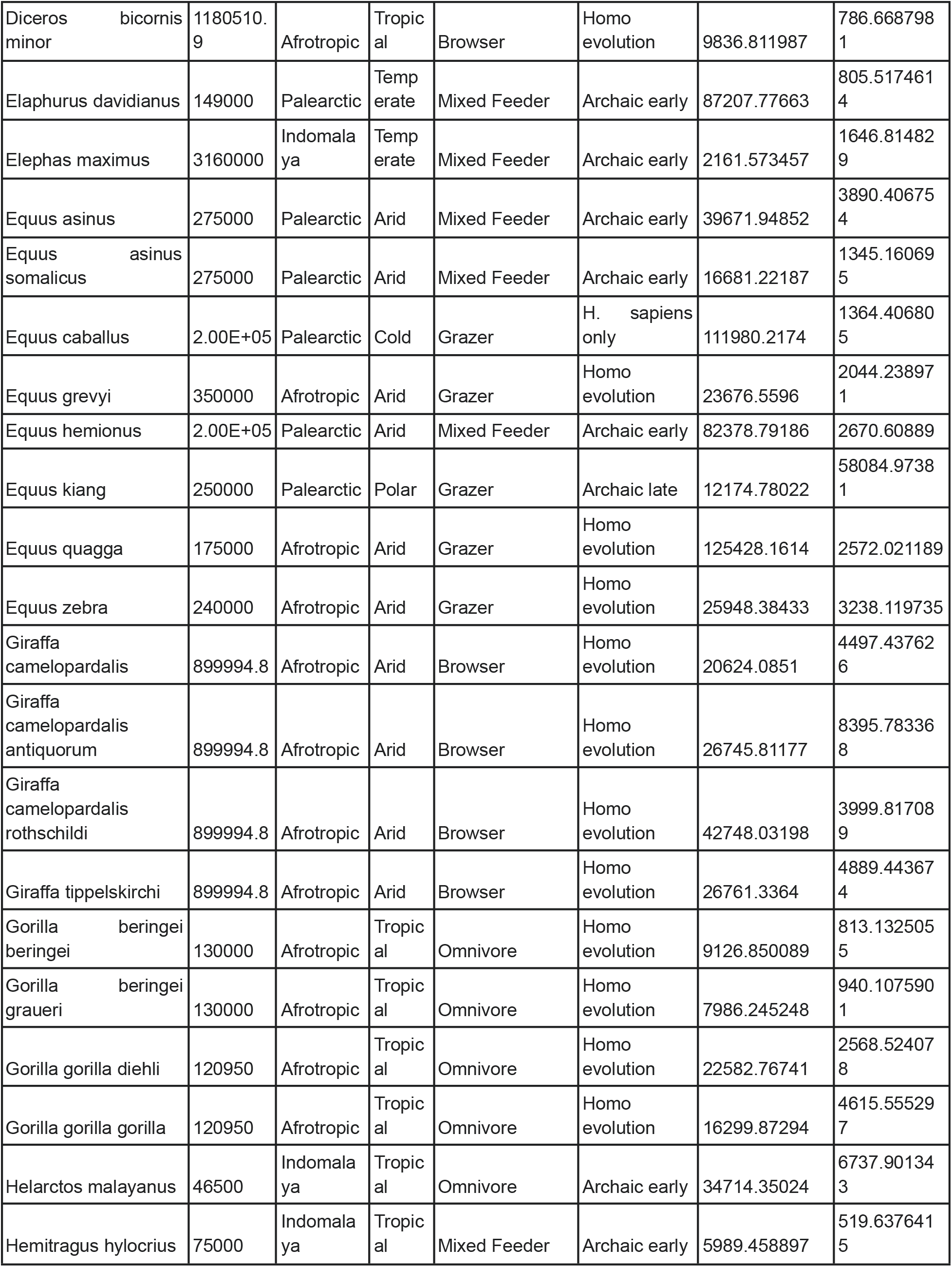
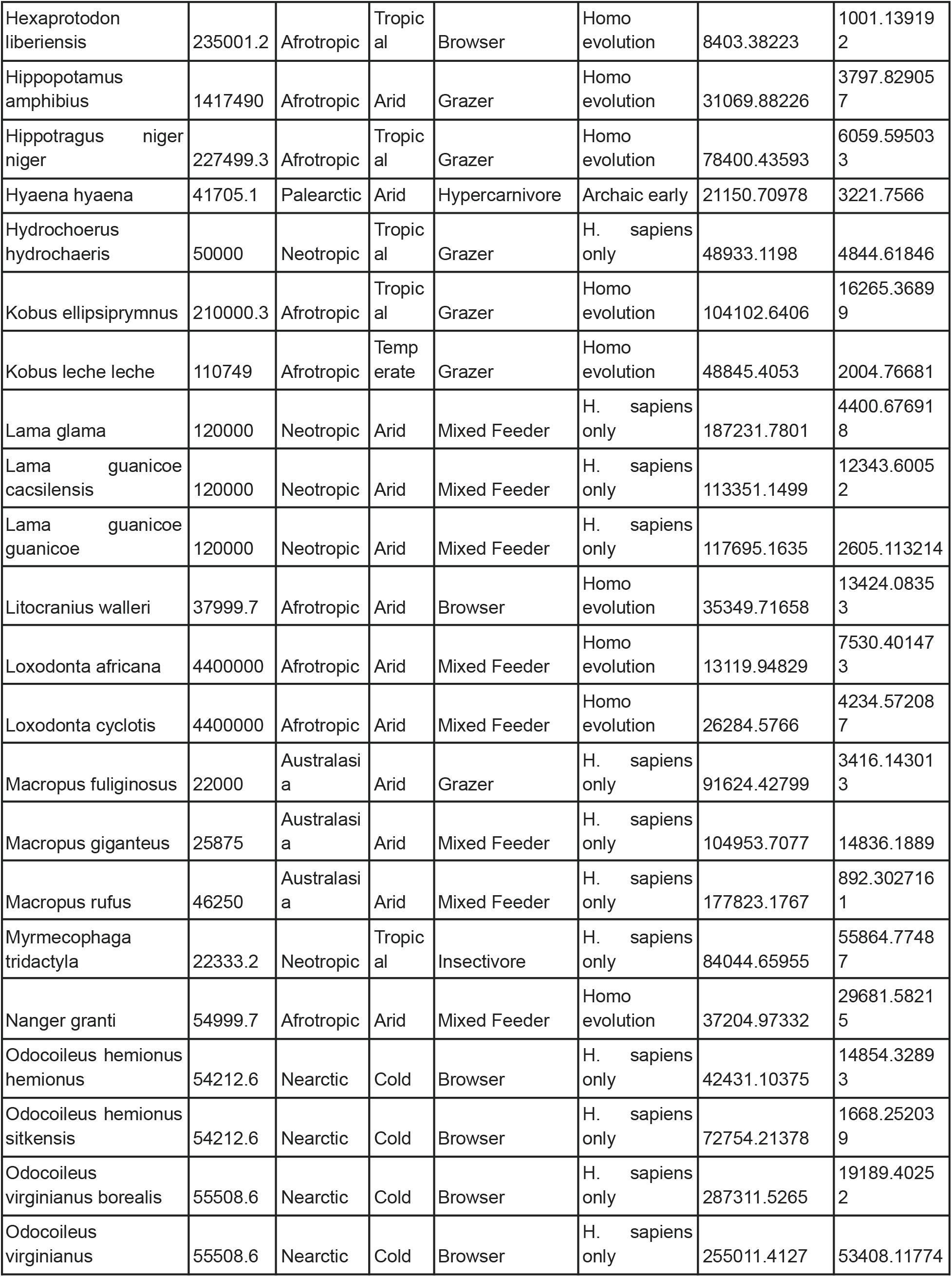
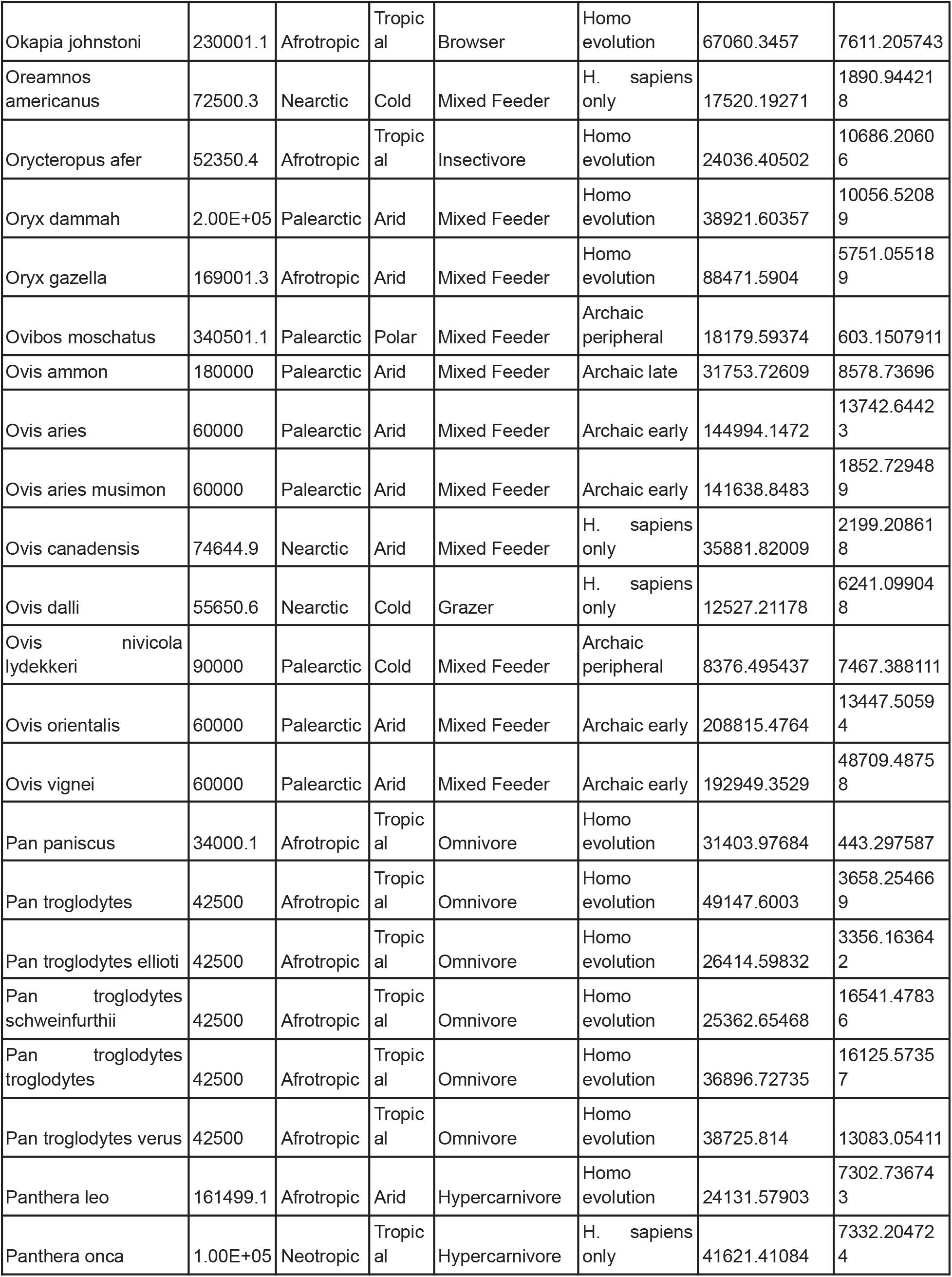
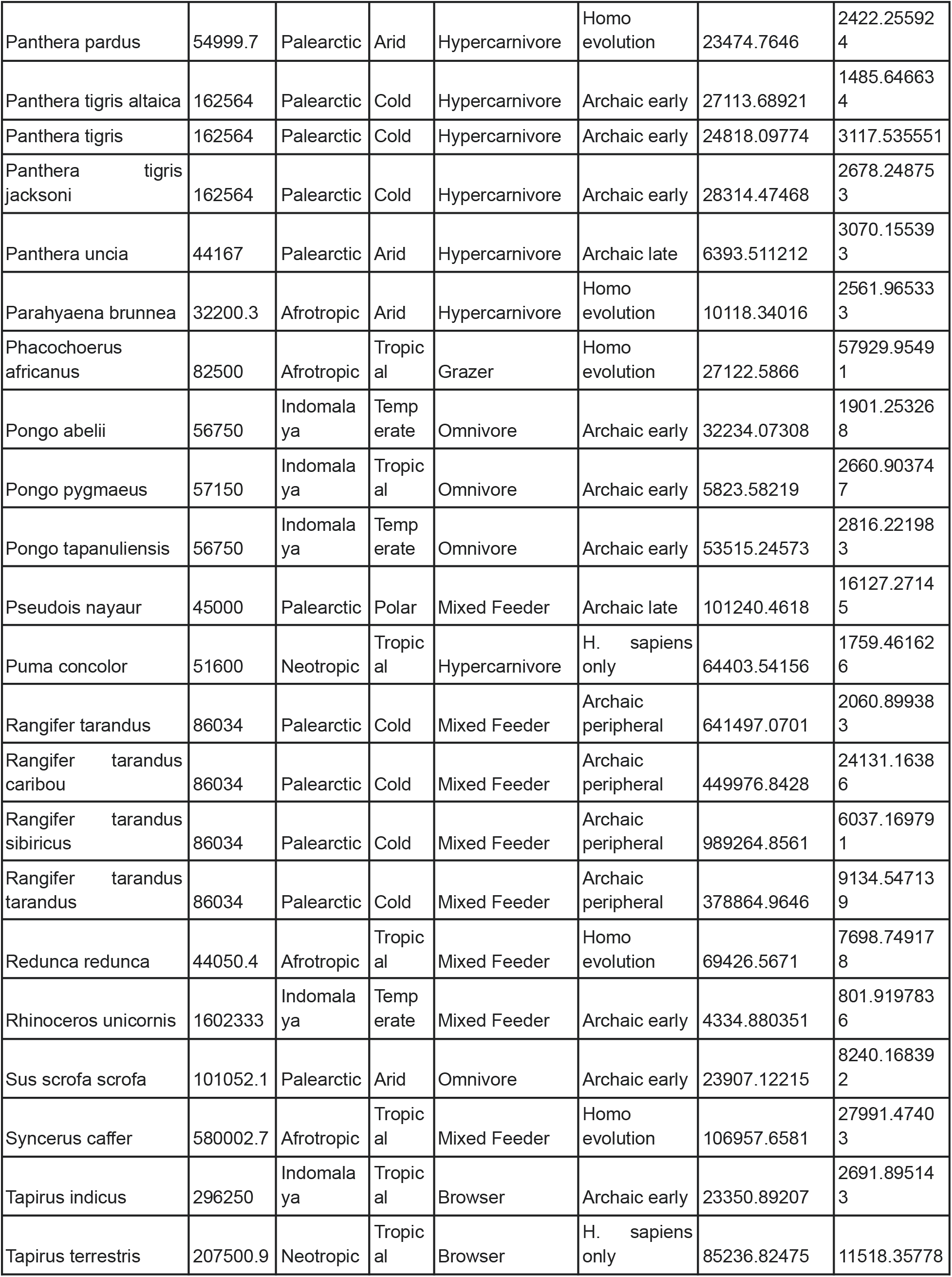
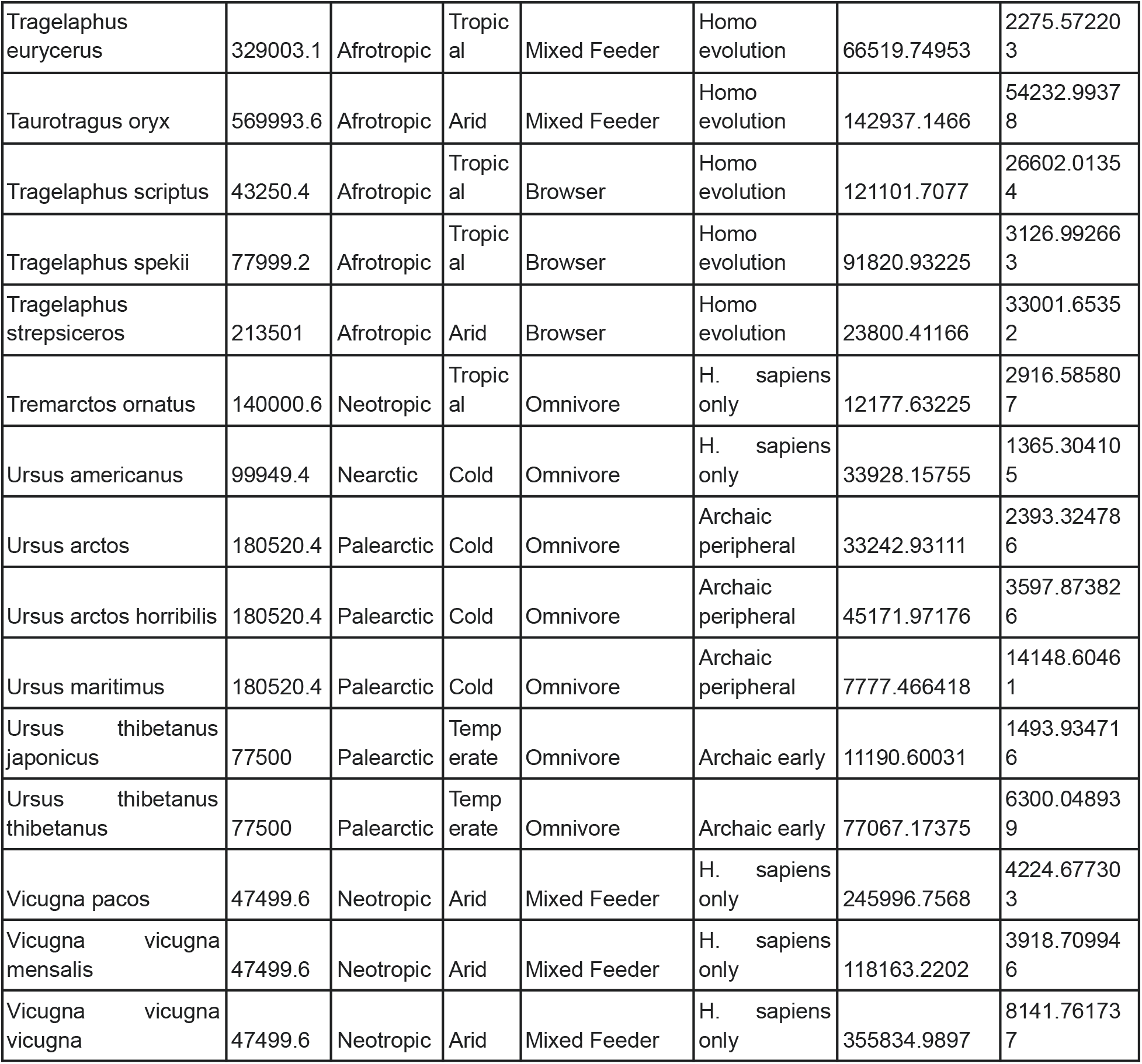
Species used in the study.

**Table S2.** Response and explanatory variables used for modelling.

| Variable | Type | Description |
| --- | --- | --- |
| $M$ | Response | Per generation mutation rate |
| $N_e$ | Response, explanatory | Average effective population size across the focal time interval |
| $D$ | Response | Decline severity |
| $N_c$ | Response | Average census population size across the focal time interval |
| $G$ | Explanatory | Generation time |
| $m$ | Explanatory | Adult mass |
| $t$ | Explanatory | Mid time-point of focal time interval |
| $t^{\text{MIN}}$ | Explanatory | Mid time-point of time interval when a species achieved the lowest population size |
| $t^{\text{MAX}}$ | Explanatory | Mid time-point of time interval when a species achieved the highest population size |
| $N_e^{\text{MAX}}$ | Explanatory | Highest past effective population size achieved by a species |
| $T$ | Explanatory | Average temperature of the focal time interval |
| $\Delta T$ | Explanatory | Difference in temperature between focal and preceding time interval |
| $L$ | Explanatory | Average temperature of the preceding time interval (temperature lag) |
| $p_H$ | Explanatory | Probability of human presence |
| $I_H$ | Explanatory | Indicator for overlap of time interval with human arrival range |

**Table S3.** Climate-based predictive models of population size.

| Name | Description |
| --- | --- |
| $\text{lin}T$ | Linear effect of temperature |
| $\text{quad}T$ | Quadratic effect of temperature |
| $\text{lin}T + L$ | Linear effect of temperature and temperature lag |
| $\text{quad}T + L$ | Quadratic effect of temperature and temperature lag |

**Table S4.** Human arrival ranges.

| Ecological realm | $a_i$ | $a_u$ |
| --- | --- | --- |
| Afrotropics | 130 kya | 200 kya |
| Australasia | 44 kya | 65 kya |
| Indomalaya | 44 kya | 73 kya |
| Nearctic | 12 kya | 20 kya |
| Neotropics | 8 kya | 16 kya |
| Paleartic | 40 kya | 95 kya |

**Table S5.** Example for the calculation of the probability of human presence *p*_*H*_.

| $a_i$ | $a_u$ | $t_i$ | $t_u$ | $p_H$ |
| --- | --- | --- | --- | --- |
| 40 kya | 95 kya | 100 kya | 125 kya | 0 |
| 40 kya | 95 kya | 75 kya | 100 kya | 0.36 |
| 40 kya | 95 kya | 50 kya | 75 kya | 0.81 |
| 40 kya | 95 kya | 25 kya | 50 kya | 1 |
| 40 kya | 95 kya | 0 kya | 25 kya | 1 |

**Table S6.** Climate-based, human-based and combined explanatory models of population size.

| Abbreviation | Class | Type |
| --- | --- | --- |
| $linT$ | Temperature only | Linear effect of temperature |
| $quadT$ | Temperature only | Quadratic effect of temperature |
| $linT + L$ | Temperature only | Linear effect of temperature and temperature lag |
| $quadT + L$ | Temperature only | Quadratic effect of temperature and temperature lag |
| $pH$ | Human only | Linear effect of probability of human presence |
| $linH$ | Human only | Linear effect of human impact post arrival |
| $expH$ | Human only | Exponential effect of human impact post arrival |
| $logH$ | Human only | Logistic effect of human impact post arrival |
| $linT + pH$ | Combined | Combination of above effects |
| $linT + linH$ | Combined | Combination of above effects |
| $linT + expH$ | Combined | Combination of above effects |
| $linT + logH$ | Combined | Combination of above effects |
| $quadT + pH$ | Combined | Combination of above effects |
| $quadT + linH$ | Combined | Combination of above effects |
| $quadT + expH$ | Combined | Combination of above effects |
| $quadT + logH$ | Combined | Combination of above effects |
| $linT + L + pH$ | Combined | Combination of above effects |
| $linT + L + linH$ | Combined | Combination of above effects |
| $linT + L + expH$ | Combined | Combination of above effects |
| $linT + L + logH$ | Combined | Combination of above effects |
| $quadT + L + pH$ | Combined | Combination of above effects |
| $quadT + L + linH$ | Combined | Combination of above effects |
| $quadT + L + expH$ | Combined | Combination of above effects |
| $quadT + L + logH$ | Combined | Combination of above effects |

### Supplementary text 1: Statistical modelling of population size

#### Mutation rate and generation time model (Figure S3)

The conversion of the PSMC output to effective population sizes and time intervals in years, requires knowledge of the per generation mutation rates and generation times. While generation times are easily obtained from literature, mutation rates are generally not available for the majority of species. However, we can use the known relationship between mutation rates and generation times in mammals to predict the mutation rate for species where these data are missing. To fit this model, we used the empirically estimated mutation rates and average parental ages (generation times) from 61 sequenced mammalian families (Bergeron *et al*., 2021; unpublished). The models is represented as follows:

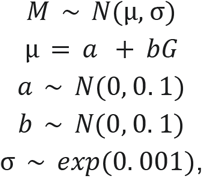

where *M* is the observed per generation mutation rate in a mammalian family and *G* is generation time. The mutation rate *M*, as well as prior distributions for the intercept (*a*) and slope (*b*) of the relationship are assumed to be normally (*N*) distributed, while the model error σ is assumed exponentially distributed (*exp*). We fit the model with the *G* predictor either on a natural scale or log-transformed. Both models have similar predictive accuracy (Figure S3), as demonstrated by the similarity of their leave-one-out cross validation log-score - 1125.94 ± 6.34 and 1124.24 ± 5.62 for the model with and without the log-transformation of *G*, respectively. However, as the validation log-score is on average higher for the model with the log-transformed *G* predictor, we use this model to predict mutation rates for species where these data are unavailable. Specifically, we use the posterior distributions of the *a* and *b* coefficients to estimate the *M* distributions for each species, and use the medians of these distributions when transforming the PSMC output.

### Category-based model across the full time span (Figure 1B)

Here, we model the change in effective population size (*N*_*e*_) of megafauna species as a function of time (*t*) while taking into account a hierarchical component that groups species into discrete categories (prior to inference, both the *N*_*e*_ and *t* values were log_10_-transformed):

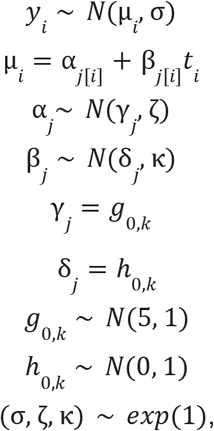

where *y*_*i*_ is an *N*_*e*_ value at a specific time in the past *t*_*i*_. The variable *y*_*i*_ is modelled as normally distributed with mean *μ*_*i*_ and standard deviation *σ*. The second line describes the global linear regression model of *N*_*e*_ as explained by *t*_*i*_, used to infer species-specific slopes (*α*_*j*_) and intercepts (*β*_*j*_), where the subscript *j* indicates a specific megafauna species. Species-specific slopes and intercepts were both modelled as response variables of a nested linear regression representing category-specific effects on the coefficients of the global model. Specifically, subscript *k* indicates a category that groups a number of species together. For example, a categorization of species into ecological realms groups species into 6 categories (Afrotropic, Australasia, Indomalaya, Nearctic, Neotropic and Palearctic species). We also considered three other species categorizations with respect to biome (5 categories: arid, cold, polar, temperate, tropical), trophic guild (6 categories: browser, grazer, hypercarnivore, insectivore, mixed feeder, omnivore) and human biogeography (5 categories: archaic early, archaic late, archaic peripheral, *H. sapiens* only, *Homo* evolution).

### Mass-based model across the full time span (Figure 1C)

Here, we model the change in effective population size (*N*_*e*_) of megafauna species as a function of time (*t*) and species’ adult mass. Prior to inference, both the *N*_*e*_ and *t* values were log_10_-transformed. The model allows for varying intercepts and slopes across species, and incorporates the effect of species adult mass as follows:

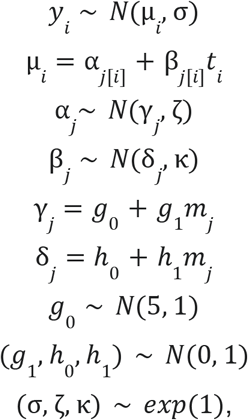

where the global regression model is identical to the previous model, but species-specific slopes and intercepts were both modelled as response variables of a nested linear regression with adult mass as an explanatory variable (*m*_*j*_). A normal distribution with mean *γ*_*j*_ (*δ*_*j*_) and standard deviation ζ (*κ*) was assumed for the species-specific slope (intercept) of the nested models. The coefficients of the nested models (*g*_{0,1}_ and *h*_{0,1}_) were subscripted with 0 for intercepts and 1 for slopes, and assigned normal prior distributions. Standard deviations of variables (σ, ζ, *κ*) were assigned an exponentially distributed prior.

### Decline severity model across the full time span (Figure 1F)

We define decline severity as

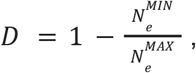

where *N*_*e*_ ^MIN^ and *N*_*e*_ ^MAX^ are the minimum and maximum effective population sizes experienced by a species during the full time interval for which we have *N*_*e*_ estimates, respectively. The Bayesian framework of the generalised linear model is as follows:

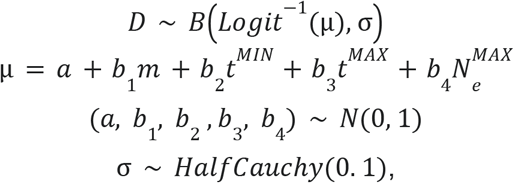

where the response variable *D* is assumed to be Beta-distributed (*B*) with mean value *μ* and standard deviation *σ*, and a logit link function. The predictors of *D* are species’ adult mass (*m*), time when a species achieved the lowest population size (*t*^MIN^), time when a species achieved the highest population size (*t*^MAX^) and the highest past population size achieved by a species (*N*_*e*_ ^MAX^), with *b*_*1*_, *b*_*2*_, *b*_*3*_, *b*_*4*_ as the corresponding coefficients and *a* as the intercept of the regression model. The predictive parameters were transformed to a scale between 0 and 1, representing the minimum and maximum value observed across species, respectively. This was done to simplify computation and achieve comparability between the estimated *b* coefficients. We assume a normally distributed prior (*N*) for the intercept and coefficients, and a Half-Cauchy (*HalfCauchy*) prior for *ϱ*.

### Climate-based predictive models (Figure 2)

Climate-based models assess the relationship between climatic variables and population size for the period between the present and 742,419 years ago (the time span for which estimates of the climatic variables are available). Here, we use *N*_*e*_ estimates from time intervals older than 100,000 years for fitting the models, while *N*_*e*_ values between the present and 100,000 years ago were predicted using the fitted models, and compared to the observed *N*_*e*_ estimates. Prior to prediction, we discretize this time frame into four equally-sized 25,000-year time windows to account for between-species differences in sizes of time windows and facilitate the comparison of model performance between time points. The goal of this modelling approach is to assess the level at which the relationship between climate fluctuations and population size prior to 100,000 years ago, can explain population size fluctuations of the recent past.

We modelled the relationship between temperature and population size using two different models. Firstly, we implemented a basic linear regression

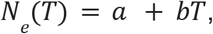

where *a* and *b* represent the intercept and slope of the relationship, while *T* is the average temperature over the focal time interval for which we have an estimate of population size *N*_*e*_. While simple, this model does not necessarily reflect biological reality due to the assumption of a linearity between population size and temperature.

To implement a more biologically realistic model, we assume that each species has an optimal temperature (*T*_*opt*_) at which *N*_*e*_ is maximised. As temperature deviates from *T*_*opt*_ in either direction, we expect a decrease in population size. Such a relationship can be described by a quadratic function

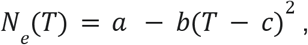

with the requirement that *b*<0. Taking the first derivative and setting the function to 0, it can be shown that the maximum of this function is *c, i*.*e. T*_*opt*_*=c*.

In addition to considering temperature of the focal interval, we also consider a temperature lag parameter *L*, defined as the average temperature of the preceding time interval.

The Bayesian frameworks are as follows:

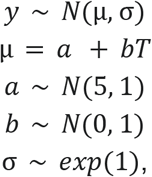

for the linear model (*linT*);

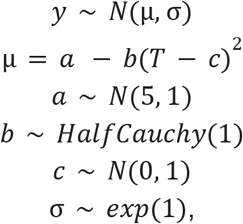

for the quadratic model (*quadT*);

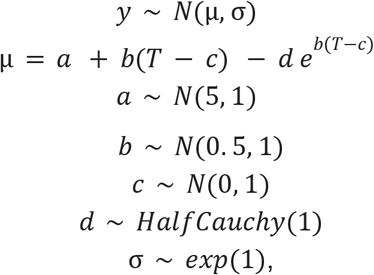

for the linear model with lag (*linT* + *L*);

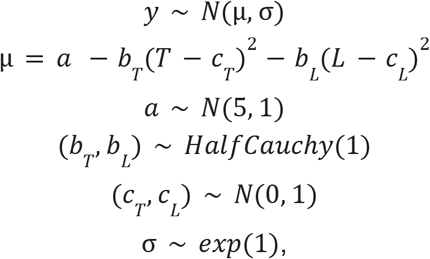

for the quadratic model with lag (*quadT* + *L*). Priors are defined as normally (*N*), exponentially (*exp*) or Half-Cauchy (*HalfCauchy*) distributed. All model parameters were estimated for each species separately apart from the model error *σ*, which is a pooled estimate across species.

### Climate and human-based explanatory models (Figure 3)

Here, we are interested in the explanatory power of climate and human impact on past population sizes of megafauna. To model the impact of climate we consider the linear and quadratic models from the previous section, in combination with human impact. The first human impact parameter we consider is the probability of human presence (*p*_*H*_), which was constructed in the following way. For each species, we consider the human arrival range based on the ecological realm of that species, and assign a value of *p*_*H*_ between 0 and 1 to each time window for which we have an estimate of the species population size. Specifically, for time windows prior to the human arrival range, *p*_*H*_ is assigned the value of 0. Windows that overlap or postcede the human arrival range are assigned a value larger than 0, depending on the span of the human arrival range and overlap of this range with the focal time window

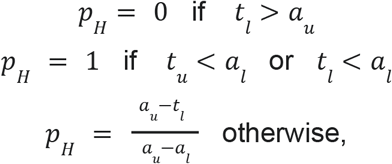

where *t*_l_ and *t*_u_ are the lower and upper bounds of the focal time window, respectively, and *a*_l_ and *a*_u_ are the lower and upper bounds of the human arrival range. Arrival time ranges for each ecological realm are given in Table S4. For the Afrotropical realm, we used the time range between 130 and 200 kya, as the time span of *H. sapiens* establishment throughout sub-Saharan Africa, while the times from Andermann et al. (2020)^16^ were taken for the other realms.

The *p*_*H*_ parameter can be thought of as cumulative human impact over time that reaches its maximum value of 1 in the time window that overlaps the lower bound of the human arrival range, and maintains this value throughout subsequent windows, towards present time. In that way, it is a conservative estimate of human impact, which, in reality, continued to increase post human arrival. Supplementary Table S5 shows an example calculation of *p*_*H*_ for five consecutive 25-ky time windows given a human arrival range between 40 and 95 kya.

The Bayesian frameworks for these models are represented as:

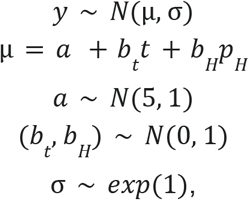

for the linear temperature and linear human impact model (*linT* + *pH*), and

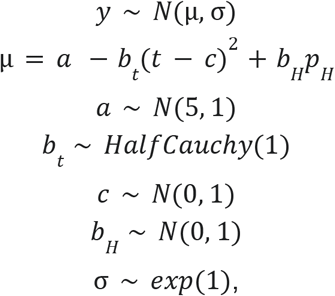

for the quadratic temperature and linear human impact model (*quadT* + *pH*). For comparison, we also run the models with only the temperature or only the human predictor, as well as models with the temperature lag parameter *L*.

We also consider a second type of human impact model, where humans are expected to start affecting megafauna population size at some point after their earliest arrival date (*a*_u_) to the ecological realm (Table S4), while prior to human arrival, we assume a constant population size. Such a model can be written as

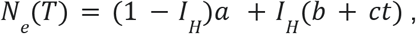

where *I*_*H*_ takes the value of 1 or 0, depending on whether or not there is overlap between human arrival range and the focal time window, respectively. Additionally, *I*_*H*_ takes the value of 1 for all windows that postcede human arrival. Further, *a* is the constant population size prior to human arrival, and the expression (*b + ct*) describes the time-dependent linear population size change following human arrival. We also consider two models with a non-linear effect post-arrival. Firstly, we consider a model with exponential impact

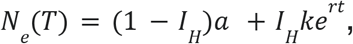

with *I*_*H*_ as in the previous model, *k* as the initial population size post arrival and *r* as the rate of population size change following human arrival. Secondly, we consider a logistic impact model

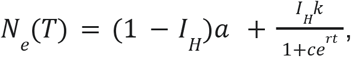

with *I*_*H*_ as in the previous model and the expression 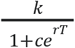 describing the logistic change in megafauna population size following human arrival. Specifically, *k* and c are constants determining the intercept of the logistic expression and *r* is the rate of population size change.

The Bayesian frameworks for these models are represented as:

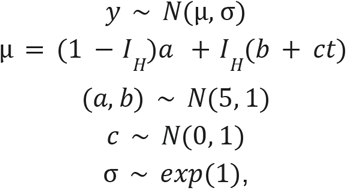

for the linear human impact model (*linH*),

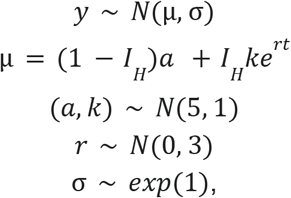

for the exponential human impact model (*expH*), and

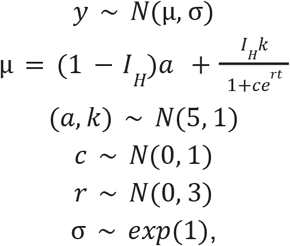

for the logistic human impact model (*logH*). Priors are defined as normally (*N*) or exponentially (*exp*) distributed. We also ran these models in combination with linear or quadratic temperature predictors. In total, we tested and compared 24 models (Table S6, Figure S2).

### Estimation of total megafauna census sizes (Figure 4C)

To estimate census sizes (*N*_*c*_) of megafauna for different time periods we utilise the positive relationship between effective and census population sizes. Specifically, we fit a linear model for the dependence of IUCN *N*_*c*_ estimates (*y*) on Holocene *N*_*e*_ estimates

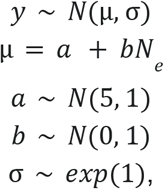

assuming a normal distribution (*N*) for the *y* parameter and the priors of the intercept (*a*) and slope (*b*) of the relationship, and an exponential (*exp*) prior for the model error σ. Furthermore, both *N*_*c*_ and *N*_*e*_ estimates were log_10_-transformed prior to model fitting, and only species with *N*_*e*_/*N*_*c*_ < 1 were used for fitting (Figure S4). The fitted model was used to infer posterior distributions of *N*_*c*_ values for each species for both the Holocene and Eemian period (based on the corresponding *N*_*e*_ values; Table S1). Each posterior distribution was randomly sampled 1,000 times to create posterior sample distributions for total megafauna census size, biomass and metabolic input for each of the two periods. Posterior sample distributions for the current, present-day period were generated in a similar way, by sampling posterior distributions of the Holocene period and then multiplying them by a scaling factor

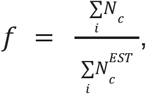

where 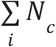 is the sum of IUCN census sizes across all species (including species with *N*_*e*_/*N*_*c*_ > 1) for which this estimate is available, while 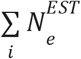 is the sum of the medians of the posterior distributions of census sizes across the corresponding species, estimated for the Holocene period. The scaling factor *f* (= 0.64) reflects the difference in current IUCN census sizes and model-predicted census sizes for the Holocene. The difference between the two sums used to calculate *f* comes from the fact that the predictive model is based only on species with *N*_*e*_/*N*_*c*_ < 1, thus considering only less severely bottlenecked species. Consequently, the posterior prediction of Holocene *N*_*c*_ for species with *N*_*e*_/*N*_*c*_ > 1 is higher than the observed IUCN values. In effect, the model predicts the Holocene census sizes that would be expected if severe bottlenecks did not occur. Arguably, this is also a more realistic scenario for the Eemian period where *N*_*c*_ values were predicted using the same model. Therefore, to obtain more realistic *N*_*c*_ values for the current period, *f* serves as a correction factor for *N*_*c*_ values estimated for the Holocene, as it reflects the average reduction between model-predicted *N*_*c*_ values, that are estimated under the assumption of lower bottleneck severities, and the observed IUCN *N*_*c*_ values, that include estimates due to severe population bottlenecks.

